# Cytosine methylation within marine sediment microbial communities: potential epigenetic adaptation to the environment

**DOI:** 10.1101/167189

**Authors:** Ian M. Rambo, Adam Marsh, Jennifer F. Biddle

## Abstract

Marine sediments harbor a vast amount of Earth’s microbial biomass, yet little is understood regarding how cells subsist in this low-energy, presumably slow-growth environment. Cells in marine sediments may require additional methods for genetic regulation, such as epigenetic modification via DNA methylation. We investigated this potential phenomenon within a shallow estuary sediment core spanning 100 years of age across its depth. Here we provide evidence of dynamic community m5-cytosine methylation within estuarine sediment metagenomes using a methylation-sensitive Illumina assay. The methylation states of individual CpG sites were reconstructed and quantified across three depths within the sediment core. A total of 6254 CpG sites were aligned for direct comparison of methylation states between samples, with 4235 sites mapped to taxa and genes. Our results demonstrate the presence of differential methylation within environmental CpG sites across an age/depth gradient of sediment. We show that epigenetic modification can be detected within complex environmental communities. The change in methylation state of environmentally relevant genes across depths may indicate a dynamic role of DNA methylation in biogeochemical processes.

## Introduction

Marine sediments are some of the largest reservoirs of microbial biomass on Earth (Whitman *et al*. 1998; Kallmeyer *et al*. 2012), and describing the relationships between community structure, activity, and ecosystem function in these habitats remains a challenge (Fuhrman 2009). The majority of sedimentary bacteria and archaea are unable to be successfully cultured in a laboratory setting, and if they are able to be cultivated, they likely do not exist in physiological states representative of those found within their natural habitats (Hoehler and Jørgensen, 2013). Next-generation sequencing technologies enable researchers to overcome the constraints of cultivation by directly analyzing environmental DNA and RNA. These technologies are employed in subsurface microbiology to provide information regarding community dynamics and ecological roles (Hua *et al*. 2014), classify rare or uncultured species (Albertsen *et al*. 2013; Seitz *et al*. 2016), and describe potential microbial activity (Orsi *et al*. 2013).

Determining the drivers that govern microbial activity in the subsurface is key to understanding the relationships between these communities and their environments. Models of the marine subsurface suggest that biomass turnover rates are on the scale of thousands of years and that many marine subsurface cells should be sporulated due to the low availability of energy (Lomstein *et al*. 2012), yet metagenomic analyses of deep-sea sediment communities exhibit low observed frequencies of endospore-specific genes (Kawai *et al*. 2015). While isolates obtained from the deep biosphere are phylogenetically similar to members of surface communities (Russell *et al*. 2016; Inagaki *et al*. 2015), cells adapted to the subsurface possibly suspend certain life processes though other functional strategies to subsist at low levels of activity. Epigenetic mechanisms offer potential microbial survival strategies within low-energy sediment, allowing for cell maintenance and acclimation to environmental stressors (Bird 2002; Casadesús and Low 2006; Low and Casadesús 2008).

DNA methylation is a conserved epigenetic modifier in prokaryotes whose roles include defense against invading foreign DNA and gene regulation (Kumar and Rao 2012; Wion and Casadesus 2006; Low *et al*. 2001; Brunet *et al*. 2011), and involves the addition of a methyl group via a DNA methyltransferase (MTase) to either the carbon 5 position of a cytosine (resulting in 5-methylcytosine (m5C)), the nitrogen 4 position of a cytosine (resulting in N4-methylcytosine (m4C)), or the nitrogen 6 position of an adenine (resulting in N6-methyladenine (m6A)) within a specific nucleotide target sequence (Ratel *et al*. 2006). These modified bases comprise an organism’s methylome, and are generally formed by two different MTase activities.

The two major forms of MTase activity are “maintenance” and “de novo” methylation. Maintenance methylation (MM) provides cells with a means of propagating DNA methylation patterns across generations. Daughter DNA strands with methylated parent strands are modified by a maintenance MTase after replication (Bird, 2002). Similarly, non-methylated parent strands normally produce non-methylated daughter strands. Unlike MM, which propagates existing methylation patterns, de novo methylation adds methyl groups to previously unmethylated bases (Kuhlmann *et al*. 2005).

While DNA methylation is an integral part of restriction-modification (RM) systems involved in the recognition of self vs. non-self DNA for cellular defense, growing evidence indicates that prokaryotes utilize both adenine and cytosine methylation as a means of regulating gene expression (Reisenauer and Shapiro 2002; Srikhanta *et al*. 2005; Srikhanta *et al*. 2009; Collier 2009; Brunet *et al*. 2011; Low and Casadesús 2008; Marinus and Casadesus 2009; Løbner-Olesen *et al.* 2005; Wion and Casadesus 2006; Low *et al*. 2001; Kahramanoglou *et al*. 2012; Gonzalez *et al*. 2014; Blow *et al*. 2016). RNA polymerase, transcription factors and binding proteins are able to recognize the methylated states of modified bases within target sites, and this discrimination of differentially methylated DNA acts as a method for determining which genes are transcribed at specific stages in the cell cycle (Low and Casadesús 2008; Gonzalez *et al*. 2014; Collier 2009).

Compared to an organism’s genome which generally remains static, the modified bases of the methylome exist in a dynamic system exhibiting plasticity outside of binary “methylated” or “non-methylated” states (Ichida *et al.*, 2007; Chernov *et al.*, 2015). A system of genetic “switches” (Hernday *et al*. 2004) regulated by dynamic DNA methylation could be a viable mechanism for both long-term and short-term transcriptional silencing for microbes inhabiting marine sediments. To better understand the prevalence and behavior of DNA methylation within marine sediment microbe communities, we utilized an Illumina sequencing-based assay to identify dynamic shifts in CpG methylation within sediment metagenomes from the Broadkill River estuary system. We opted to utilize this assay due to the presence of cytosine methylation at CpG sites in prokaryotes and the anticipation of a heterogeneous, dynamic community. Since adenine methylation is considered to be more widespread than cytosine methylation in prokaryotes (Ratel *et al*. 2006), this choice of motif also serves to reduce signal saturation in this mixed community. To the best of our knowledge, this is the first report on DNA methylation within metagenomic sequence data, and is the first to utilize this method of CpG methylation analysis in an environmental application.

## Materials and Methods

### Core collection

Sediment cores were sampled from the Oyster Rocks site of the Broadkill River, Milton, DE, USA (38.802161, −75.20299) at low tide in July 2012 and 2014. The 2012 core was sectioned into 3 cm sections and immediately frozen at −80°C for subsequent processing and DNA extraction. The sediment collection from 2012 was depleted to extract sufficient DNA for sequencing. Three cores were extracted from the same site in 2014 ~5 m from the riverbank: a 32 cm radionuclide dating core (R), and 25 cm (S) and 30 cm (L) cores for pore water ion chromatography, methane flame ionization gas chromatography, and porosity measurements. Cores L and S were sliced into 3 cm depth samples and immediately frozen at −80 °C, while Core R was immediately processed.

### Radionuclide dating

Core R was sectioned into 1 cm thick intervals from 0–10 cm, and 2 cm thick intervals from 10−32 cm. Samples were weighed, dried at 60 °C for 48 hours, reweighed, and transferred to a 25 °C desiccation chamber for storage until further processing. Dried samples were crushed with a mortar and pestle, and ground into a fine powder with an IKA Werke M20 mill (IKA Werke, Staufen, Germany). Powdered samples were transferred to 60 ml plastic jars and compressed at 3.4x10^3^ kPa with a manual hydraulic press. Radionuclide counting of compressed samples was performed for 24 hours on a Canberra Instruments Low Energy Germanium Detector (Canberra Industries, Meriden, CT, USA). Levels of ^7^Be (t_1/2_ = 53.22 days), ^210^Pb (t_1/2_ = 22.20 years), and ^137^Cs (t_1/2_ = 30.17 years) activity were measured by gamma spectroscopy of the 478, 46.5, and 662 kEV photopeaks, respectively (Igarashi *et al*. 1998; Cutshall *et al*. 1983; Wallbrink *et al*. 2002).

### Porewater ion chromatography

Porewater was extracted from 50 mL sediment samples by centrifugation at 13,000 G for 30 minutes. Porewater ions were measured with a Metrohm 850 Professional ion chromatograph (Metrohm, Herisau, Switzerland). Dilutions were measured to determine a standard curve. Samples were diluted to ensure signal within the standard curve.

### Methane flame ionization gas chromatography

Methane concentrations were determined for Core L and S subsamples (volume = 305 cm^3^) extracted from each core slice using a 5 ml syringe whose top had been removed with a sterile razor blade. Core subsamples were transferred into 20 mL amber glass vials, and 1 mL 1 M NaOH was added to each vial to halt microbial activity. Vials were crimped, shaken, and stored for 10 days at 25°C. A standard curve was calculated from 500, 1000, and 5000 ppm standards. Mean headspace methane concentrations were determined by running 100 µL gas extractions in triplicate via flame ionization gas chromatography using a 5890 Series II gas chromatograph equipped with a flame ionization detector (Hewlett-Packard, Palo Alto, California, USA).

### Metagenome library preparation and sequencing

Metagenome libraries were prepared from the 2012 sediment core sections. Genomic DNA (gDNA) was extracted from 0.5 g of sediment with a MoBio PowerSoil (MoBio, Valencia, CA) kit per the manufacturer’s protocol. A 10 µg aliquot of purified gDNA was digested with the methylation-sensitive restriction endonuclease HpaII, which cleaves at the unmodified internal cytosine of a 5’-CCGG-3’ motif. Digested DNA was cleaned with a QIAquick PCR purification kit (Qiagen, Hilden, Germany), sheared to a median size of 300 bp using a Covaris focused-ultrasonicator (Covaris, Woburn, MA, USA), and cleaned again with QIAquick. Digested extracts were immediately transferred to −20°C until library preparation. Illumina libraries were prepared using the NEBNext Ultra Library Prep Kit for Illumina (New England BioLabs, Ipswich, MA, USA) and sequenced with an Illumina Hi-Seq 2500 (Illumina, San Diego, California, USA) at the Delaware Genomics and Biotechnology Institute (Newark, DE, USA). Single-read sequencing was performed for all samples, with 150-cycle sequencing for the 3–l6 cm and 12–15 cm samples, and 50-cycle sequencing for the 24–27 cm sample. All sequence reads are deposited in GenBank under the study PRJEB11699.

### 16S rRNA gene amplicon sequencing and analysis

DNA was extracted, purified, and digested using the previously described method for Illumina libraries. Purified DNA was quantified and tested for successful PCR reactions for the bacterial 16S rRNA gene. Amplicon library preparation and sequencing of 16S rRNA genes were performed by Molecular Research, LP (Clearwater, Texas, USA).

Analysis of 16S rRNA gene sequences was performed with QIIME 1.8.0 (Caporaso *et al.*, 2010). Dereplication, abundance sorting, and discarding reads less than 2 bp was performed with the USEARCH7 algorithm (Edgar, 2013). Chimeras were filtered with UCHIME (Edgar *et al.*, 2011) using the RDP Gold Classifier training database v9 (Cole *et al.*, 2014). Operational taxonomic unit (OTU) picking was performed at 97% similarity with UCLUST (Edgar, 2010). Non-chimeric sequences were chosen as the representative set of sequences for taxonomic assignment and alignment. Taxonomic assignments were performed with UCLUST (Edgar 2010) using the Greengenes V13.8 database for 97% OTUs (DeSantis *et al*. 2006). OTU tables were rarefied from 2000 to 9500 sequences per sample by steps of 100, with 10 iterations performed at each step.

### Metagenome assembly and annotation

Metagenome sequence reads were trimmed to 51 bp and quality controlled to only include those with Phred nucleotide confidence scores greater than or equal to 95%. Quality-controlled reads were assembled in IDBA (Peng *et al.*, 2010) with parameters –mink 18 –maxk 36 –step 2 –similar 0.97 –min_count 2 (Table S1). Phylogenetic classification of IDBA-assembled contigs was performed with PhymmBL (Brady and Salzberg, 2011) and Kraken (Wood and Salzberg, 2014). A PhymmBL identity confidence score threshold of 65% was imposed to designate higher-confidence Order-level assignments. Comparative taxonomic classifications were performed with Kraken (Wood and Salzberg, 2014) using the standard database comprised of complete RefSeq bacterial, archaeal, viral, and fungal genomes. Contigs assigned to viral or fungal genomes in Kraken were removed from downstream analyses. Marker gene annotation of filtered contigs was performed with Phylosift (Darling *et al.*, 2014).

Open reading frame (ORF) prediction was performed in six reading frames with MetaGene (Noguchi *et al*. 2006). ORFs were annotated for KEGG Orthology (KO) families (Kanehisa *et al.*, 2016) in HMMER 3.0 (Eddy, 2011) using the Functional Ontology Assignments for Metagenomes (FOAM) database (Prestat *et al.*, 2014) and an e-value acceptance threshold of 1e-4. In the case of multiple KO assignments per contig, the result with the best e-value and bitscore was chosen to represent that contig. Contigs that did not receive a protein annotation from these software were aligned with BLASTX (Altschul *et al.*, 1997) against the NCBI non-redundant protein database and scored with the BLOSUM62 substitution matrix (Henikoff and Henikoff, 1992), with a maximum expectation value of 1e-4 and a word size of 3.

### Metagenome CpG methylation quantification and statistical analysis

CpG methylation was calculated using a custom bioinformatic pipeline and software platform (Genome Profiling LLC; Marsh and Pasqualone 2014). Platform workflow is managed down a decision tree from assembled contigs, and performs the following tasks: 1) isolation of informative target contigs, 2) sequence compression to reduce complexity, 3) contig re-assembly and mapping to reference metagenomes, 4) CpG quantification for m5C site distributions (% methylation across all gDNA copies in the sample), and 5) methylation profiling comparison between samples based on quantitative statistics. The methylation score metrics recovered from assembled contigs are based on independent characteristics of DNA fragmentation via HpaII restriction digest and random shearing.

Computational reconstruction of CpG methylation is based on a null selection model, where the distribution of m5C modifications at any single CpG site is expected to be 50% in a large population of cells (genome copies) for CpG sites that are non-functional or silent. Where CpG methylation status is important for cellular fitness and thus there is a selection force pushing the m5C distributions away from a 50:50 equilibrium, these scoring algorithms are focused on quantifying this departure from the null expectation, and the degree of departure measured is proportional to overall methylation status of that CpG site among all the cellular genome copies being sampled.

Statistical analyses of methylation scores were performed using R statistical package. Modalities were tested with Hartigans’ dip test for unimodality. Methylation score bootstrap standard errors (SE) and coefficients of variation (CV) were estimated (n = 10,000). Score variances were tested with a Brown-Forsythe Levene-type test. Two-tailed Jonckheere-Terpstra trend tests were performed with 10,000-permutation reference distributions.

## Results

### Sediment properties

Radionuclide dating constraints show that the Oyster Rocks site is comprised of a top layer of recently deposited tidally mixed or bioturbated sediment (~ 4 cm, sediment age < 106 days) situated above older sediment established 50–100+ years ago (Figure S1). Sulfate concentrations were more varied between 0–3 cm and 3–6 cm (Figure S1 A) for Core L, but concentrations were higher in deeper samples from 6–9 cm to 27–30 cm. Methane concentrations of Core L were shown to increase with depth, with higher variance between 0–12 cm and lower variance from 15–30 cm (Figure S1 B). Porosity for Core R was shown to be far lower within older sediments (Figure S1 C). The higher variability of SO_4_^−2^ and CH_4_ in more recently deposited sediments could be due to tidal forcing. While cores from separate years were used to generate sequence and geochemical data, the ages of all sediments are consistent, in that the shallowest sequenced sample is less than a year old and the deeper sequenced samples are significantly older (50+ years). In general, the deepest samples are anoxic, with a transition occurring past 9 cm depth. This follows the trends established in earlier sampling in this area (Cheng 2013), showing that generalities can be drawn over time.

### 16S rRNA gene analysis

The diversity of 16S rRNA genes was generally higher at 3–6 cm than the 12–15 cm and 24–27 cm samples (Figure S2). The 12–15 cm and 24–27 cm samples had similar profiles for rarefied Chao1 diversity and observed OTU counts. These deeper samples also had a higher presence of Dehalococcoidetes and sulfate-reducing Deltaproteobacteria, as well as Marine Crenarchaoetal Group and Marine Hydrothermal Vent Group archaea (Figure S3). OTUs were clearly shared between the three depths, and corresponding abundance changes suggest that known anaerobic taxa were more abundant at depth.

### Metagenome taxonomic composition and function

The most abundant taxonomic classes present in metagenomic and 16S rRNA gene data across all depths were the Actinobacteria, Bacilli, Clostridia, Deinococci, and α-β-δ-γ-proteobacteria (Figure 1, S4). The sediment community transitions to a mostly anaerobic environment with depth based on taxonomic and functional annotations. Metagenome contigs with both higher-scoring PhymmBL annotations and Kraken annotations further indicate a prevalence of anaerobic taxa within the 12–15 and 24–27 cm samples. The presence of methanogenic archaea (Methanomicrobiales and Methanosarcinales) and anaerobic Dehalococcoidia and Deltaproteobacteria within the 12–15 and 24–27 cm samples (Figure 1, S4) suggest these communities support anaerobic lifestyles (Oremland and Polcin, 1982). KO annotations suggest that deeper communities have the potential for anaerobic metabolism (Figure 2), as greater abundances of genes involved in sulfate reduction (formate dehydrogenase, adenylyl sulfate kinase, NADH dehydrogenase, and heterodisulfide reductase) and methanogenesis (trimethylamine corrinoid protein co-methyltransferase) were present within the 24–27 cm sample.

**Figure 1:**
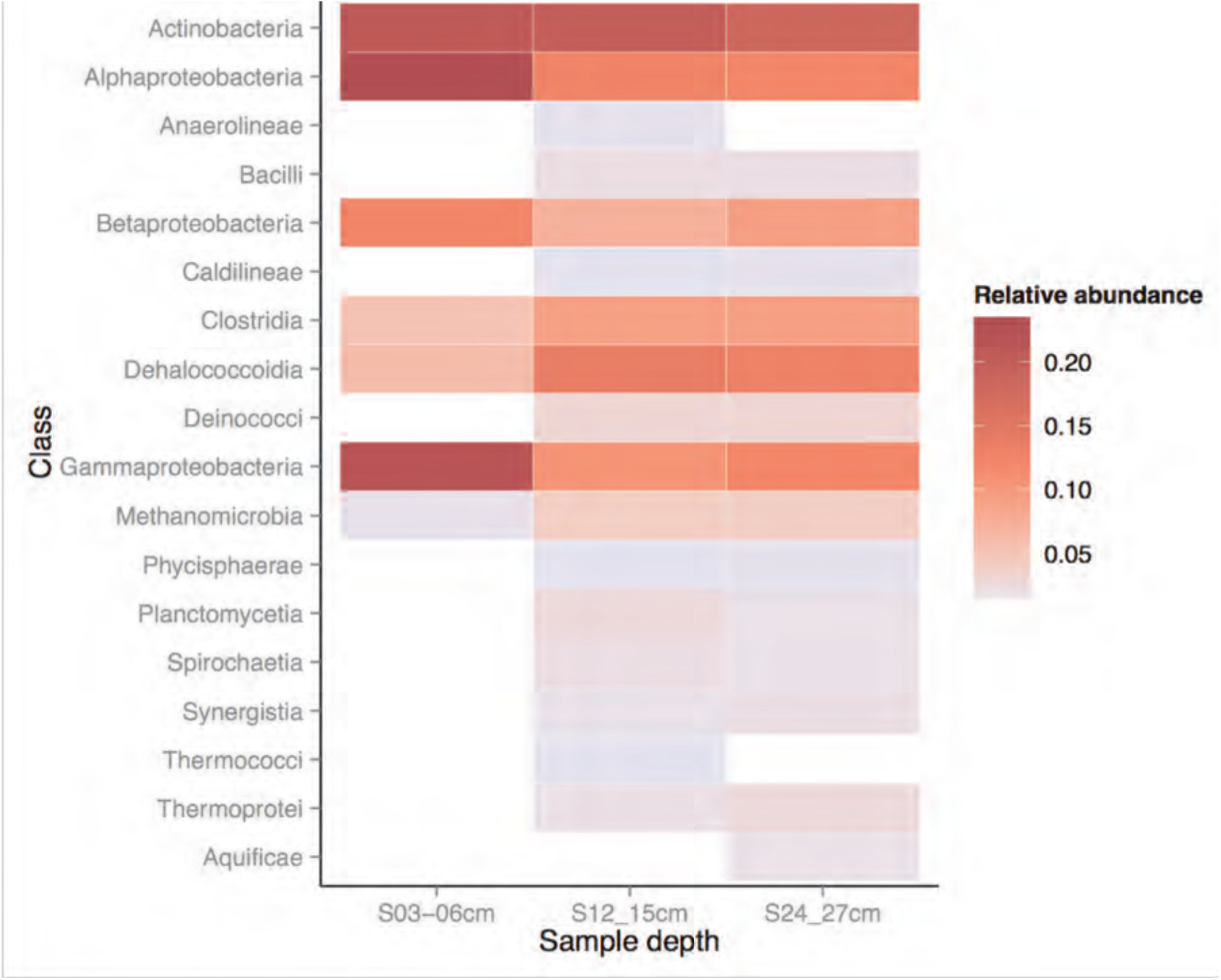
Taxonomic relative abundances >= 0.05% of metagenome contigs with both Kraken and higher-confidence PhymmBL class assignments present in two or more samples. PhymmBL annotations were paired with Kraken annotations to select against potential false positives annotated by PhymmBL. These annotations suggest a presence of Actinomycetales at all depths, and an increased abundance of anaerobic classes (Clostridia, Dehalococcoidia). Methanogenic archaea (Methanosarcinales) are present in the 12–15 cm and 24–27 cm samples.

**Figure 2:**
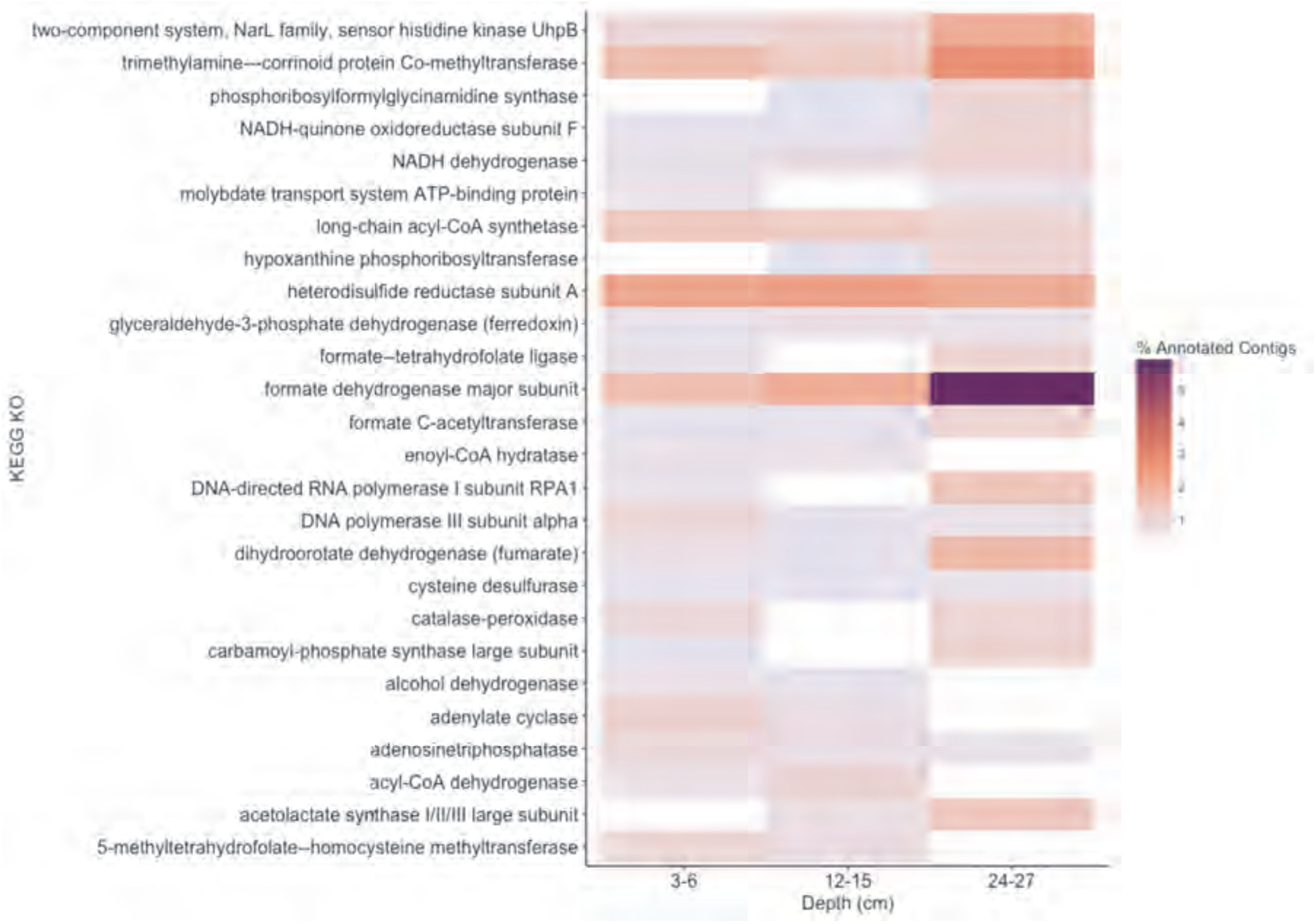
Abundance of KEGG Orthology functional assignments across sample depths. KEGG Orthology annotation results show that Oyster Rocks sediment communities at 12–15 cm and 24–27 cm have higher enzymatic potential for anaerobic metabolism.

### Metagenome CpG methylation

From these metagenome data, we assessed the methylation states of CpG sites. A total of 6254 CpG sites that could be directly compared between all three samples were mapped to 3743 contigs (4.33% of all three unprocessed IDBA assemblies). Differential methylation states were observed in 1173 sites, while the remaining 5081 had equivalent methylation states. Of these CpG sites, 4235 (67.7%) were identified within contigs receiving higher-confidence PhymmBL Order classifications.

The methylation shift behaviors of individual CpG sites are varied and highly dependent upon their original states. Community-wide methylation distributions showed higher proportions of CpG sites that remain in highly methylated states from 3–6 cm to 12–15 cm (Figure 3, A; Figure 4 A), and can be associated with MM. A gradual overall loss in methylation and transition into more equilibrated, binary states of high and low methylation was seen at 12–15 cm (Figure 3, B) and 24–27 cm (Figure 3, C). An apparent increase of methylation losses ranging from ~25–50% was shown to account for this (Figure 4, B), with higher numbers of sites shifting from ~80–90% methylated states to hemimethylated and non-methylated states (Figure 5, B). A greater number of CpG sites experienced shifts from non-methylated states to fully methylated states when transitioning from the 12–15 cm to 24–27 cm, potentially indicating *de novo* methylation of these sites. It should be noted that many CpG sites were shown to remain in non-methylated states between 12–15 cm and 24–27 cm.

**Figure 3:**
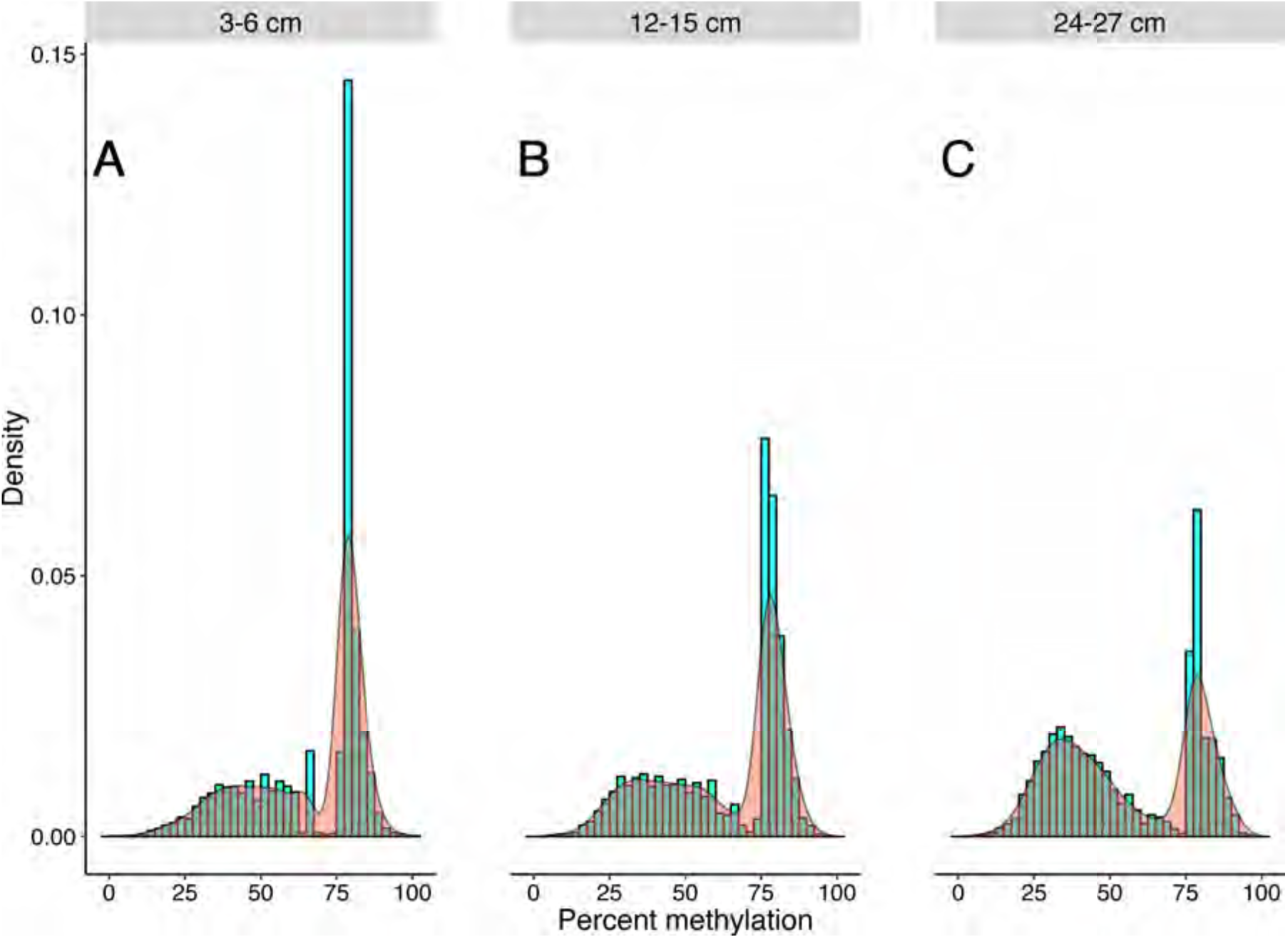
Community methylation load distributions. Histograms and kernel density overlays represent overall methylation levels of CpG sites that are shared across all three samples. A greater number of recovered CpG sites were highly methylated at 3–6 cm (**A**), resulting in a greater methylation load at this depth. However, a trend of decreasing overall methylation was seen at 12–15 cm (**B**) and 24–27 cm (**C**), with more sites experiencing transitions from highly methylated to non-methylated or hemimethylated states, or persisting in non-methylated states.

**Figure 4:**
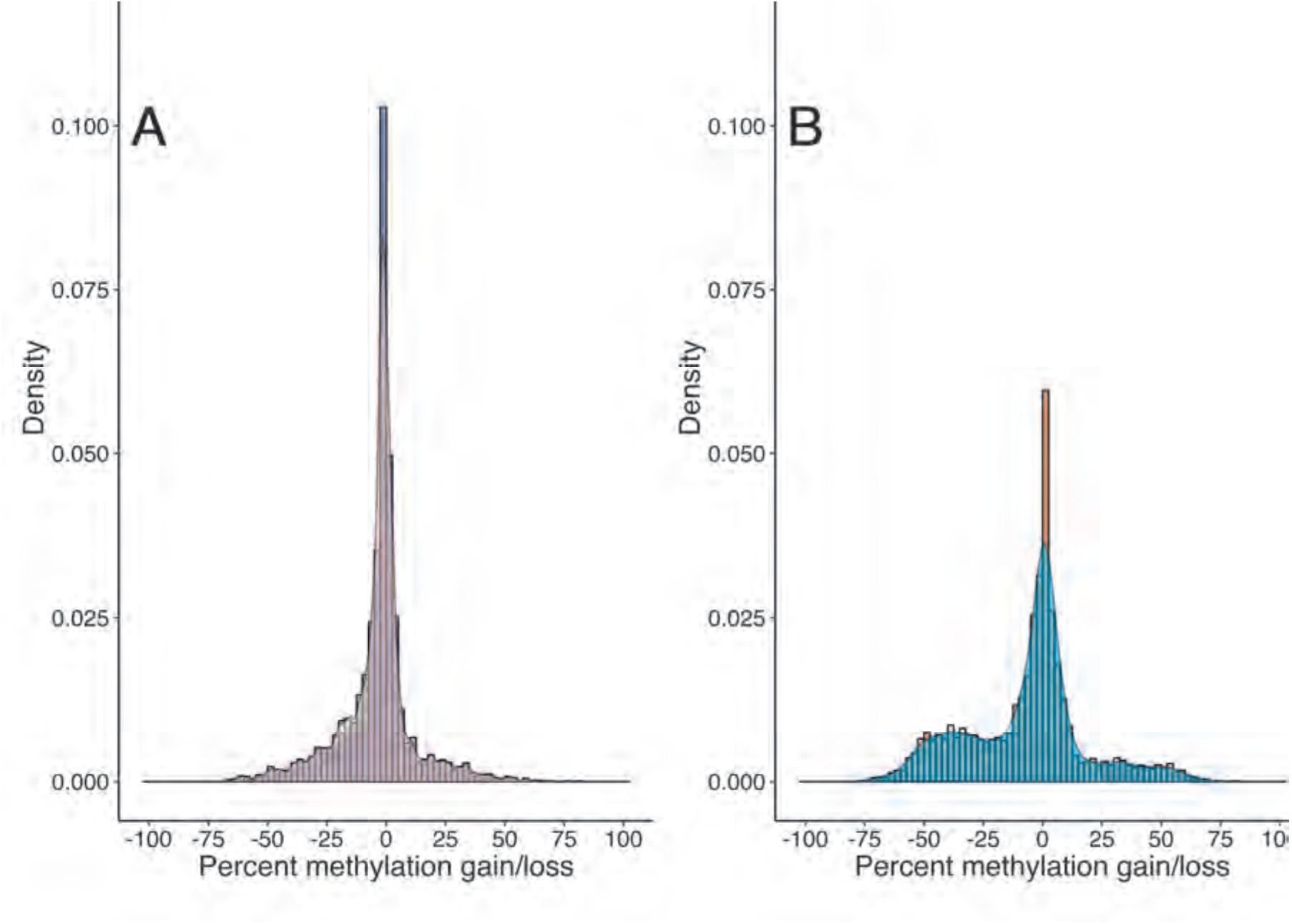
CpG methylation gains and losses from (**A**) 3–6 cm to 12–15 cm and (**B**) 12–15 cm to 24–27 cm. Histograms and kernel density overlays are representative of the densities of methylation shifts for individual CpG sites. Sites represented in (**A**) are the same sites represented in (**B**). Shifts range from −100 (total methylation loss) to +100 (total methylation gain). A significant number of CpG sites remained at equivalent methylation states from 3–6 cm to 12–15 cm, yet there is an apparent increase in methylation losses ranging from ~25% to ~50% from 12–15 cm to 24–27 cm.

**Figure 5:**
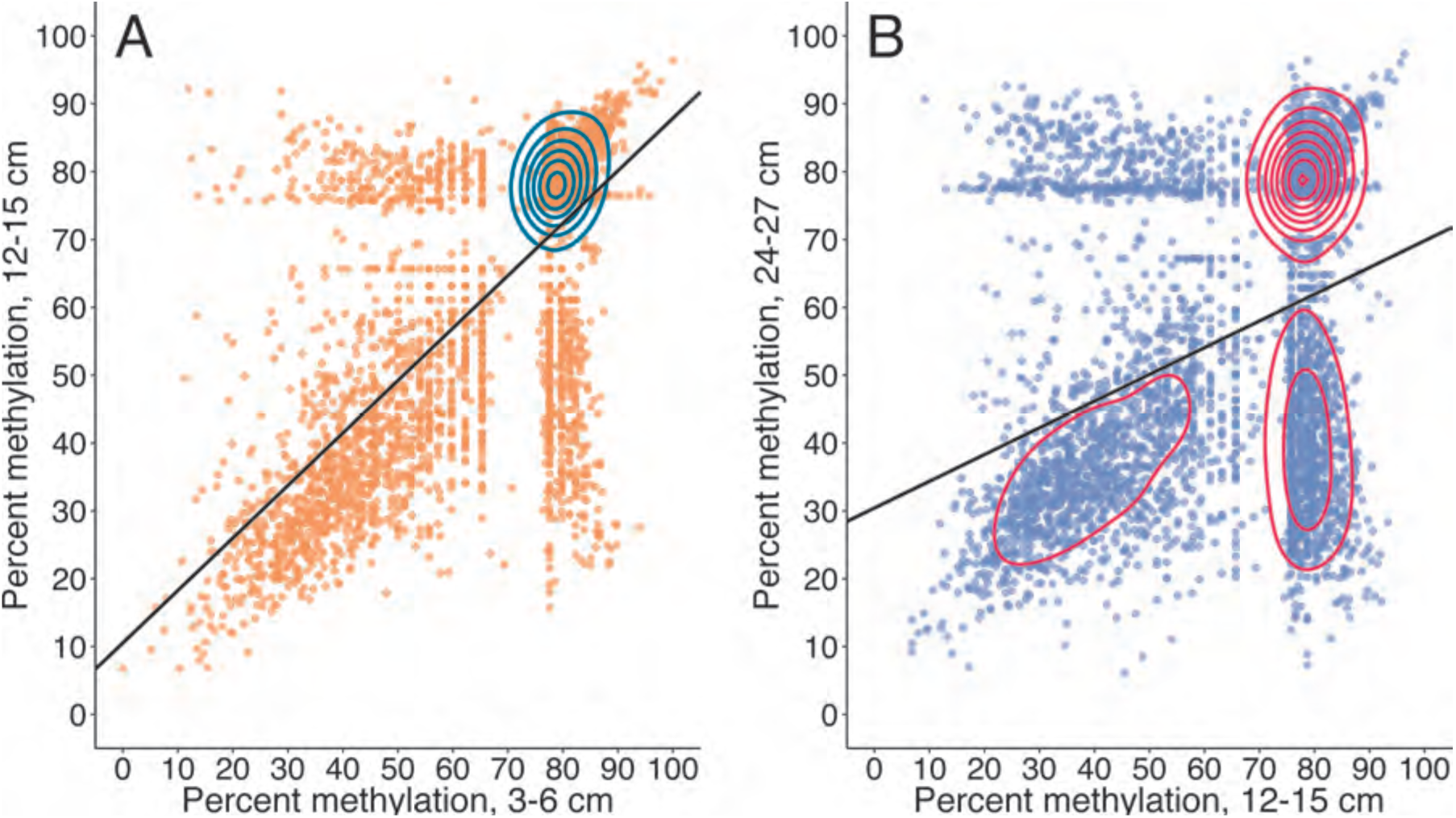
Total CpG methylation shifts from 3–6 cm to 12–15 cm (**A**) and 12–15 cm to 24–27 cm (**B**). Plots are representative of metagenome-wide methylation loads. Each point is a recovered CpG site whose quantified methylation states are comparable across all three samples. The CpG sites represented in (**A**) are the same as those represented in (**B**). Changes in a CpG site’s methylation profile can be traced from a shallower sample (x-axis) to a deeper sample (y-axis). A greater number of sites remain in highly methylated states from 3–6 cm to 12–15 cm. These same CpG sites experience a general trend of methylation loss from the mid sample to the deepest sample. An increased number of CpG sites with methylation loads ~80% at 12–15 cm undergo methylation losses ranging from 20–60% upon transitioning to 24–27 cm.

The methylation dynamics of individual CpG sites were analyzed for taxa with higher numbers of recovered sites. An overall trend of increasing methylation score standard error (SE) and coefficient of variation (CV) with depth was seen in all analyzed phyla (Table S2). There is a general trend of decreasing CV for methylation scores with depth, and this is influenced by an overall trend towards bimodal score distributions. Hartigans’ dip test results support a non-unimodal distribution of methylation scores for analyzed phyla (Table S3), verifying mixed methylation profiles. Brown-Forsythe tests suggest that CpG score variances across depths were unequal for 70% of analyzed phyla (p < 0.05), supporting the presence of mixed methylation profiles and dynamic shifts in methylation states (Table S4). Jonckheere-Terpstra trend test results show that community methylation scores decrease overall with depth (*p* = 2e-4). Methylation scores for the majority of phyla exhibit decreasing trends with depth (Table S4; Figure 3; Figure 5).

Only 35 CpG sites were mapped to contigs receiving KEGG Orthology annotations. Of these 35 CpG sites, chitinase gene annotations were recovered for 14 comparable sites that could be traced back to six contigs with higher-confidence PhymmBL classifications (Figure 6). A total of 12 quantifiable sites exhibiting differential methylation states within the same gene were identified for Actinomycetales and Thermoanaerobacterales.

**Figure 6:**
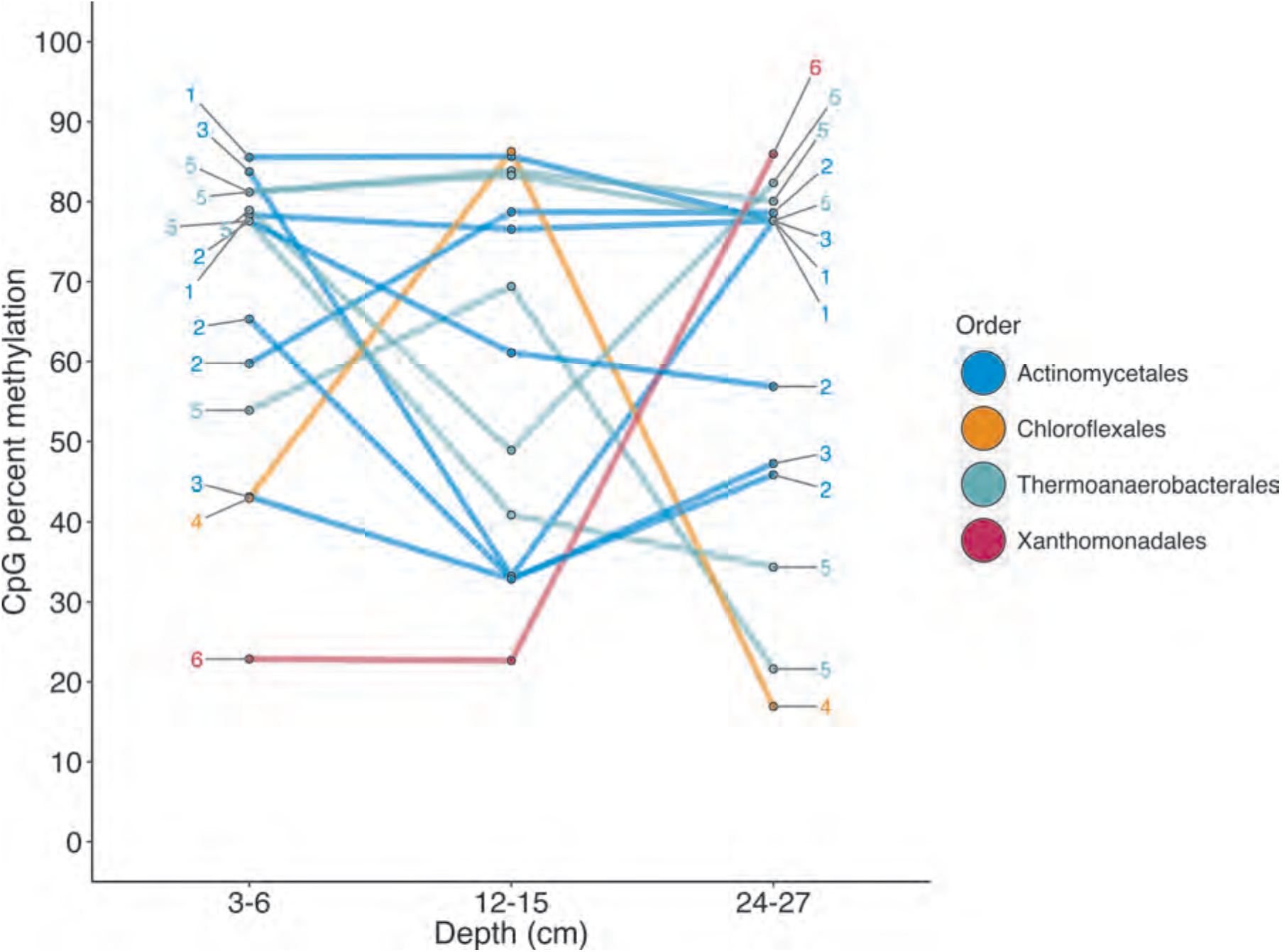
Chitinase CpG methylation dynamics. Quantifiable CpG sites were recovered for six contigs mapped to four orders. Connected points are representative of changes in the methylation state of a single CpG site for each sample. Numbers denote contigs to which these CpG sites are mapped. Multiple CpG sites exhibiting differential methylation within the same gene were recovered for the Actinobacteria and Thermoanaerobacterales. The recovered CpG site mapped to Xanthomonadales was shown to persist in a non-methylated state from 3–6 cm to 12–15 cm, but exhibited an apparent *de novo* methylation event from 12–15 cm to 24–27 cm.

Quantifiable states of 73 CpG sites were recovered for transposase genes identified by BLASTX alignments (Figure 7). Transposase CpG sites that were methylated in surface samples tended to remain in methylated states across depths, although several methylated sites undergo shifts into hemimethylated or non-methylated sites in deeper samples. CpG sites existing within the same contig tended to shift to similar methylation states from 12–15 cm to 24–27 cm.

**Figure 7:**
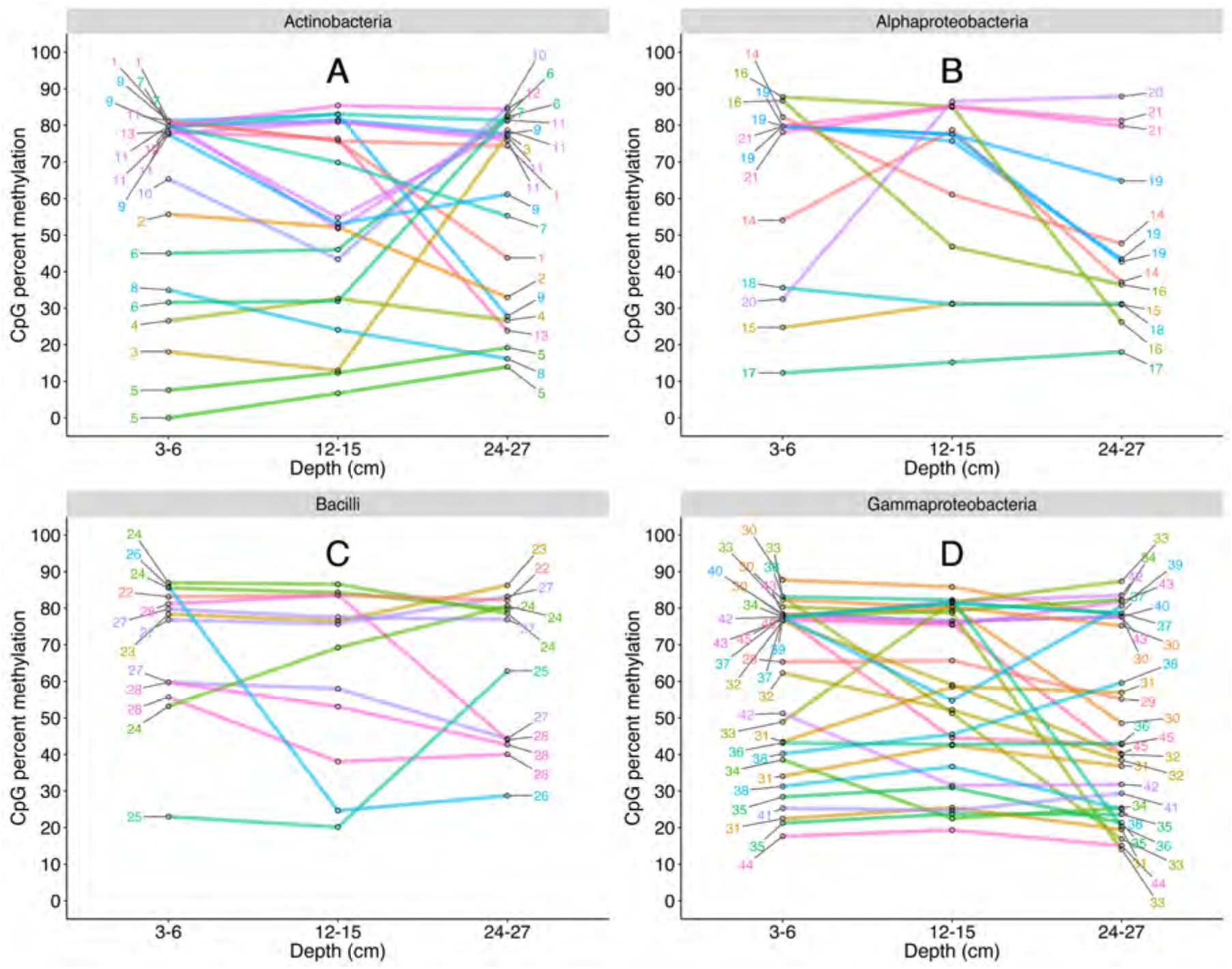
CpG site methylation shifts for transposases mapped to Actinobacteria (**A**), Alphaproteobacteria (**B**), Bacilli (**C**), and Gammaproteobacteria (**D**). Interpretation guidelines are the same as those in Figure 6. CpG sites were shown to exist in differential methylation states across depths. The methylation states of multiple CpG sites located within the same contig were shown to shift to highly similar states in several instances.

## Discussion

DNA methylation has an established association with RM systems and gene regulation within cultured prokaryotes (Casadesús and Low 2006; Blow *et al*. 2016). However, its presence and function within uncultured environmental microbes is not well understood, especially considering the extent of undiscovered taxa (Locey and Lennon, 2016). Current methylation detection protocols for NGS sequencing are improbable for sediment samples due to community complexity, high required concentration and molecular weight of DNA samples, and error rate. We provide evidence of differential m5C methylation at CpG sites within estuarine sediment communities using a restriction enzyme-based Illumina sequencing method capable of reconstructing methylation profiles from environmental DNA samples. This ability for continuous CpG methylation scoring within metagenomic DNA opens future possibilities for better understanding how microbes in the subsurface can respond to various conditions and stressors.

Sediment communities observed through both 16S rRNA gene and metagenomic gDNA sequencing do not appear to be governed highly by factors such as porosity and sediment porewater geochemistry, and this notion is supported by previous research of Broadkill River sediments (Cheng 2013) and other estuarine microbe communities (Koretsky *et al.*, 2005). However, shifts in community composition appear to be more closely related to the drastic change in sediment age suggested by radionuclide constraints.

Analysis of 16S rRNA gene amplicons suggests that the 3–6 cm sample communities exhibit greater diversity than the deeper samples (Figure S2). It is likely that surface sediments are more aerobic than deeper sediments due to regular cycling and deposition, and these higher oxygen levels could be a factor in this higher diversity. Obligate anaerobes and facultative aerobes were observed in the shallow sample as well. The 3–6 cm sample also encompasses the transition zone from young, fresh sediment to older, established sediment at 4–5 and 5–6 cm, so overlap in communities was expected. Our results clearly show increases in age from 3–6 to 12–15 cm, and the deeper depth of 24–27 cm is certainly older although our tests could not measure an exact age between 15–24 cm. Results support the presence of a drastically shifting downcore age gradient with higher anaerobic community potential at depth. Methane and porewater ion profiles are more varied within surface sediments, suggesting a bioturbated or tidally mixed region of fresh sediment in line with ^7^Be and ^210^Pb activity constraints. CpG methylation profiles recovered from these sediments were mapped to taxa and genes, and exhibit dynamic shifts in methylation state.

Of the CpG site populations assigned to taxa, 22% had < 5% methylation gains or losses between depths. Two possible explanations are presented here. First, MM can propagate methylated CpGs and result in a higher conservation of methylated states. Second, MM does not act upon non-methylated CpG sites, and as such these sites tend to remain non-methylated (Casadesús and Low, 2006). While these sites remained within generally equivalent states, 9.22% of recovered CpGs were observed to have differential methylation shifts >= 50% between depths. This is representative of the standard binary response associated with the concept of an epigenetic on/off switch (Marsh and Pasqualone 2014; Hernday *et al*. 2004). Shifts between highly methylated and fractionally methylated states suggest the presence of dynamic CpG sites that contribute to a mixed population. However, we cannot rule out the potential effect of gene or whole genome duplication on methylation scoring, as newly replicated DNA would contain fewer methylated bases and is highly dependent upon maintenance methylation (Casadesús and Low, 2006).

CpG methylation states were shown to vary for specific genes including chitinases and transposases. Chitinolytic bacteria are widely distributed in sediment environments (Bhattacharya *et al*. 2007; Souza *et al*. 2011), and are responsible for converting this insoluble source of carbon and nitrogen into a widely used form. It has been previously noted that chitin is rapidly removed from an estuary within the first 10 cm of sediment (Gooday 1990). The most abundant shifts from the 3–6 cm to 12–15 cm samples are from methylated to unmethylated states within lineages of Actinobacteria and Thermoanaerobacter, both of which can degrade chitin (Bhattacharya *et al*. 2007; Spanevello and Patel 2015). We interpret the variations in these signals to suggest regulation, and not just a signature of cellular replication, considering that methylation losses are greater upon transition to the anaerobic 12–15 cm depth. These same CpG sites are again methylated within the 24–27 cm depth as they leave the assumed zone of available chitin. While chitin was not concurrently measured, the often noted correlation between cultivable chitinolytic bacteria and chitin abundances suggests that this process is one that would not be maintained if chitin were not present (Gooday 1990). The evidence for anaerobic organisms only reducing methylation from chitinase CpG sites within the anaerobic sedimentary horizons suggests that this is methylation-based regulation of an metabolically energetically costly process. We postulate that this is an initial glimpse into how marine sediment microbes potentially utilize DNA methylation to regulate biogeochemical processes that are vital for nutrient cycling.

DNA methylation’s potential to regulate gene transcription also applies to the expression of genes involved in the transport of transposable elements, i.e. transposases. We provide evidence for differential methylation of multiple CpG sites within sediment bacterial transposase genes (Figure 7), as a lack of methylation within CpG sites or shifts into hemimethylated states could hint at the potential activity of transposases. The regulation of transposases and transposon mobility could play a role in rapid acclimation responses by influencing transcriptional activity and the ability for mobile genes to be inserted into a genome. Transposase regulation has been observed to take place via adenine methylation at GATC sites in *E. coli* (Dodson and Berg 1989; Reznikoff 1993; Roberts *et al*. 1985; Yin *et al*. 1988; Spilemann-Ryser *et al.* 1991), yet the regulatory mechanisms of one model organism do not necessarily apply to the entire bacterial domain. Horizontal gene transfer is speculated to occur in estuarine sediment microbes (Angermeyer *et al*. 2016), and extracellular transposases could play a potential role in this process as substantial numbers of horizontally-transferred genes in several bacteria species are attributed to foreign DNA such as transposons (Ochman *et al*. 2000). Due to the known influence of DNA methylation within bacterial transposons and the results of this study, we speculate that DNA methylation could act as a regulator of transposition within the subsurface.

As epigenetic research shifts from model systems towards potentially novel organisms within natural environments, there is a pressing need for the development of assays capable of detecting epigenetic signatures within environmental samples. This study provides a community-level insight into the dynamic behavior of a well-known and conserved methylation site within estuarine sediments. A benefit of this Illumina assay is that it requires less DNA than single-molecule approaches, and allows for CpG site mapping to specific taxa and genes. Future modifications tailored for metagenomic samples could pave the way for the reconstruction of dynamic methylation profiles within genomes obtained from the environment.

## Acknowledgements

This work was funded in part by the Center for Dark Biosphere Investigations (CDEBI; NSF OCE-0939564) and the University of Delaware. Thanks to Annamarie Pasqualone for sample preparation, Caitlyn Tucker and Chris Sommerfield for radionuclide expertise, Tom Hanson for ion chromatography, Rovshan Mahmudov for FIGC, and Patrick Gaffney for statistics expertise. This is CDEBI publication number XXX.

## Conflict of Interest

The software platform designed for processing DNA methylation profiles from NGS sequence data is licensed by the University of Delaware to Genome Profiling LLC, a company co-founded by Adam G. Marsh and developed with support from an Innovation Corps Grant from the National Science Foundation to AGM. Ian Rambo and Jennifer Biddle declare no financial involvement with any commercial entity.

Datasets for this project are publicly available from BCO-DMO at http://www.bcodmo.org/dataset/628253.

Supplemental Information includes figures and tables.

## References

Albertsen, M., Hugenholtz, P., Skarshewski, A., Nielsen, K. L., Tyson, G. W. and Nielsen, P. H. (2013) ‘Genome sequences of rare, uncultured bacteria obtained by differential coverage binning of multiple metagenomes.’, Nature biotechnology, 31(6), pp. 533–8. doi: 10.1038/nbt.2579.

Altschul, S. F., Madden, T. L., Schäffer, A. A., Zhang, J., Zhang, Z., Miller, W. and Lipman, D. J. (1997) ‘Gapped BLAST and PSI-BLAST: A new generation of protein database search programs’, Nucleic Acids Research, 25(17), pp. 3389–3402. doi: 10.1093/nar/25.17.3389.

Angermeyer, A., Crosby, S. C. and Huber, J. A. (2016) ‘Decoupled distance-decay patterns between dsrA and 16S rRNA genes among salt marsh sulfate-reducing bacteria’, Environmental Microbiology, 18(1), pp. 75–86. doi: 10.1111/1462-2920.12821.

Bhattacharya, D., Nagpure, A. and Gupta, R. K. (2007) ‘Bacterial chitinases: Properties and Potential.’, Critical reviews in biotechnology, 27(1), pp. 21–28. doi: 10.1080/07388550601168223.

Bird, A. (2002) ‘DNA methylation patterns and epigenetic memory.’, Genes & development, 16(1), pp. 6–21. doi: 10.1101/gad.947102.

Blow, M. J., Clark, T. A., Daum, C. G., Deutschbauer, A. M., Fomenkov, A., Fries, R., Froula, J., Kang, D. D., Malmstrom, R. R., Morgan, R. D., Posfai, J., Singh, K., Visel, A., Wetmore, K., Zhao, Z., Rubin, E. M., Korlach, J., Pennacchio, L. A. and Roberts, R. J. (2016) ‘The Epigenomic Landscape of Prokaryotes’, PLoS Genetics, 12(2), pp. 1–28. doi: 10.1371/journal.pgen.1005854.

Brady, A. and Salzberg, S. (2011) ‘PhymmBL expanded: confidence scores, custom databases, parallelization and more’, Nature methods, 8(5), pp. 2011–2013. doi: 10.1038/nmeth0511-367.

Brunet, Y. R., Bernard, C. S., Gavioli, M., Lloubès, R. and Cascales, E. (2011) ‘An epigenetic switch involving overlapping fur and DNA methylation optimizes expression of a type VI secretion gene cluster.’, PLoS genetics, 7(7), p. e1002205. doi: 10.1371/journal.pgen.1002205.

Caporaso, J. G., Kuczynski, J., Stombaugh, J., Bittinger, K., Bushman, F. D., Costello, E. K., Fierer, N., Peña, A. G., Goodrich, J. K., Gordon, J. I., Huttley, G. a, Kelley, S. T., Knights, D., Koenig, J. E., Ley, R. E., Lozupone, C. a, Mcdonald, D., Muegge, B. D., Pirrung, M., Reeder, J., Sevinsky, J. R., Turnbaugh, P. J., Walters, W. a, Widmann, J., Yatsunenko, T., Zaneveld, J. and Knight, R. (2010) ‘correspondence QIIME allows analysis of high- throughput community sequencing data Intensity normalization improves color calling in SOLiD sequencing’, Nature Publishing Group. Nature Publishing Group, 7(5), pp. 335–336. doi: 10.1038/nmeth0510-335.

Casadesús, J. and Low, D. (2006) ‘Epigenetic gene regulation in the bacterial world.’, Microbiology and molecular biology reviews: MMBR, 70(3), pp. 830–56. doi: 10.1128/MMBR.00016-06.

Cheng, B. (2013) Variations of Archaeal Communities in Sediments of Coastal Delaware. University of Delaware, Thesis.

Chernov, A. V., Reyes, L., Peterson, S. and Strongin, A. Y. (2015) ‘Depletion of CG-specific methylation in Mycoplasma hyorhinis genomic DNA after host cell invasion’, PLoS ONE, 10(11), pp. 1–14. doi: 10.1371/journal.pone.0142529.

Cole, J. R., Wang, Q., Fish, J. a., Chai, B., McGarrell, D. M., Sun, Y., Brown, C. T., Porras-Alfaro, A., Kuske, C. R. and Tiedje, J. M. (2014) ‘Ribosomal Database Project: Data and tools for high throughput rRNA analysis’, Nucleic Acids Research, 42(D1), pp. 633–642. doi: 10.1093/nar/gkt1244.

Collier, J. (2009) ‘Epigenetic regulation of the bacterial cell cycle.’, Current opinion in microbiology, 12(6), pp. 722–9. doi: 10.1016/j.mib.2009.08.005.

Darling, A. E., Jospin, G., Lowe, E., Matsen, F. a, Bik, H. M. and Eisen, J. a (2014) ‘PhyloSift: phylogenetic analysis of genomes and metagenomes.’, PeerJ, 2, p. e243. doi: 10.7717/peerj.243.

Eddy, S. R. (2011) ‘Accelerated profile HMM searches’, PLoS Computational Biology, 7(10). doi: 10.1371/journal.pcbi.1002195.

Edgar, R. C. (2010) ‘Search and clustering orders of magnitude faster than BLAST’, Bioinformatics, 26(19), pp. 2460–2461. doi: 10.1093/bioinformatics/btq461.

Edgar, R. C. (2013) ‘UPARSE: highly accurate OTU sequences from microbial amplicon reads.’, Nature Methods, 10(10), pp. 996–8. doi: 10.1038/nmeth.2604.

Edgar, R. C., Haas, B. J., Clemente, J. C., Quince, C. and Knight, R. (2011) ‘UCHIME improves sensitivity and speed of chimera detection’, Bioinformatics, 27(16), pp. 2194–2200. doi: 10.1093/bioinformatics/btr381.

Fuhrman, J. a (2009) ‘Microbial community structure and its functional implications.’, Nature, 459(7244), pp. 193–9. doi: 10.1038/nature08058.

Gonzalez, D., Kozdon, J. B., McAdams, H. H., Shapiro, L. and Collier, J. (2014) ‘The functions of DNA methylation by CcrM in Caulobacter crescentus: a global approach.’, Nucleic acids research, 42(6), pp. 3720–35. doi: 10.1093/nar/gkt1352.

Gooday, G. W. (1990) ‘The ecology of chitin degradation’, Advances in Microbial Ecology, 11, pp. 387–430.

Henikoff, S. and Henikoff, J. G. (1992) ‘Amino acid substitution matrices from protein blocks’, Proceedings of the National Academy of Sciences of the United States of America, 89(November), pp. 10915–10919.

Hernday, A., Braaten, B. A. and Low, D. A. (2004) ‘The intricate workings of a bacterial epigenetic switch’, Advances in experimental medicine and biology, 547.

Hoehler, T. M. and Jørgensen, B. B. (2013) ‘Microbial life under extreme energy limitation.’, Nature reviews. Microbiology. Nature Publishing Group, 11(2), pp. 83–94. doi: 10.1038/nrmicro2939.

Hua, Z.-S., Han, Y.-J., Chen, L.-X., Liu, J., Hu, M., Li, S.-J., Kuang, J.-L., Chain, P. S., Huang, L.-N. and Shu, W.-S. (2014) ‘Ecological roles of dominant and rare prokaryotes in acid mine drainage revealed by metagenomics and metatranscriptomics.’, The ISME journal. Nature Publishing Group, pp. 1–15. doi: 10.1038/ismej.2014.212.

Ichida, H., Matsuyama, T., Abe, T. and Koba, T. (2007) ‘DNA adenine methylation changes dramatically during establishment of symbiosis’, FEBS Journal, 274(4), pp. 951–962. doi: 10.1111/j.1742-4658.2007.05643.x.

Inagaki, F., Kubo, Y., Bowles, M. W., Heuer, V. B., Ijiri, A., Imachi, H., Ito, M., Kaneko, M., Lever, M. A., Morita, S., Morono, Y., Tanikawa, W., Bihan, M., Bowden, S. A., Elvert, M., Glombitza, C., Gross, D., Harrington, G. J., Hori, T., Li, K., Limmer, D., Murayama, M., Ohkouchi, N., Ono, S., Purkey, M., Sanada, Y., Sauvage, J., Snyder, G., Takano, Y., Tasumi, E., Terada, T., Tomaru, H., Wang, D. T. and Yamada, Y. (2015) ‘Exploring deep microbial life in coal-bearing sediment down to ~2.5 km below the ocean floor’, Science, 349(6246), pp. 420–424.

Kahramanoglou, C., Prieto, A. I., Khedkar, S., Haase, B., Gupta, A., Benes, V., Fraser, G. M., Luscombe, N. M. and Seshasayee, A. S. N. (2012) ‘Genomics of DNA cytosine methylation in Escherichia coli reveals its role in stationary phase transcription’, Nature Communications. Nature Publishing Group, 3, p. 886. doi: 10.1038/ncomms1878.

Kallmeyer, J., Pockalny, R., Adhikari, R. R., Smith, D. C. and D’Hondt, S. (2012) ‘Global distribution of microbial abundance and biomass in subseafloor sediment.’, Proceedings of the National Academy of Sciences of the United States of America, 109(40), pp. 16213–6. doi: 10.1073/pnas.1203849109.

Kanehisa, M., Sato, Y., Kawashima, M., Furumichi, M. and Tanabe, M. (2016) ‘KEGG as a reference resource for gene and protein annotation’, Nucleic Acids Research, 44(D1), pp. D457–D462. doi: 10.1093/nar/gkv1070.

Kawai, M., Uchiyama, I., Takami, H. and Inagaki, F. (2015) ‘Low frequency of endospore-specific genes in subseafloor sedimentary metagenomes’, Environmental Microbiology Reports, 7(2), pp. 341–350. doi: 10.1111/1758-2229.12254.

Koretsky, C. M., Van Cappellen, P., DiChristina, T. J., Kostka, J. E., Lowe, K. L., Moore, C. M., Roychoudhury, A. N. and Viollier, E. (2005) ‘Salt marsh pore water geochemistry does not correlate with microbial community structure’, Estuarine, Coastal and Shelf Science, 62(1–2), pp. 233–251. doi: 10.1016/j.ecss.2004.09.001.

Kuhlmann, M., Borisova, B. E., Kaller, M., Larsson, P., Stach, D., Na, J., Eichinger, L., Lyko, F., Ambros, V., Söderbom, F., Hammann, C. and Nellen, W. (2005) ‘Silencing of retrotransposons in Dictyostelium by DNA methylation and RNAi’, Nucleic Acids Research, 33(19), pp. 6405–6417. doi: 10.1093/nar/gki952.

Kumar, R. and Rao, D. N. (2012) ‘Role of DNA Methyltransferases in Epigenetic Regulation in Bacteria’, in Epigenetics: Development and Disease, pp. 81–102. doi: 10.1007/978-94-007-4525-4.

Løbner-Olesen, A., Skovgaard, O. and Marinus, M. G. (2005) ‘Dam methylation: Coordinating cellular processes’, Current Opinion in Microbiology, 8(2), pp. 154–160. doi: 10.1016/j.mib.2005.02.009.

Locey, K. J. and Lennon, J. T. (2016) ‘Scaling laws predict global microbial diversity’, Proceedings of the National Academy of Sciences, Early edit, pp. 1–6. doi: 10.7287/peerj.preprints.1451v1.

Lomstein, B. A., Langerhuus, A. T., D’Hondt, S., Jørgensen, B. B. and Spivack, A. J. (2012) ‘Endospore abundance, microbial growth and necromass turnover in deep sub-seafloor sediment.’, Nature. Nature Publishing Group, 484(7392), pp. 101–4. doi: 10.1038/nature10905.

Low, D. A., Weyand, N. J. and Mahan, M. J. (2001) ‘Roles of DNA Adenine Methylation in Regulating Bacterial Gene Expression and Virulence’, Infection and Immunity, 69(12), pp. 7197–7204. doi: 10.1128/IAI.69.12.7197.

Low, D. and Casadesús, J. (2008) ‘Clocks and switches: bacterial gene regulation by DNA adenine methylation.’, Current opinion in microbiology, 11(2), pp. 106–12. doi: 10.1016/j.mib.2008.02.012.

Marinus, M. G. and Casadesus, J. (2009) ‘Roles of DNA adenine methylation in host-pathogen interactions: Mismatch repair, transcriptional regulation, and more’, FEMS Microbiology Reviews, 33(3), pp. 488–503. doi: 10.1111/j.1574-6976.2008.00159.x.

Marsh, A. G. and Pasqualone, A. A. (2014) ‘DNA methylation and temperature stress in an Antarctic polychaete, Spiophanes tcherniai’, Frontiers in Physiology, 5(May), pp. 1–9. doi: 10.3389/fphys.2014.00173.

Noguchi, H., Park, J. and Takagi, T. (2006) ‘MetaGene: Prokaryotic gene finding from environmental genome shotgun sequences’, Nucleic Acids Research, 34(19), pp. 5623–5630. doi: 10.1093/nar/gkl723.

Ochman, H., Lawrence, J. G. and Groisman, E. A. (2000) ‘Lateral gene transfer and the nature of bacterial innovation’, Nature, 405, pp. 299–304.

Oremland, R. S. and Polcin, S. (1982) ‘Methanogenesis and sulfate reduction: competitive and noncompetitive substrates in estuarine sediments.’, Applied and Environmental Microbiology, 44(6), pp. 1270–1276. doi: 10.1016/0198-0254(83)90262-5.

Orsi, W. D., Edgcomb, V. P., Christman, G. D. and Biddle, J. F. (2013) ‘Gene expression in the deep biosphere.’, Nature. Nature Publishing Group, 499(7457), pp. 205–8. doi: 10.1038/nature12230.

Peng, Y., Leung, H. C. M., Yiu, S. M. and Chin, F. Y. L. (2010) ‘IDBA - A practical iterative De Bruijn graph De Novo assembler’, Lecture Notes in Computer Science (including subseries Lecture Notes in Artificial Intelligence and Lecture Notes in Bioinformatics), 6044 LNBI, pp. 426–440. doi: 10.1007/978-3-642-12683-3_28.

Prestat, E., David, M. M., Hultman, J., Taş, N., Lamendella, R., Dvornik, J., Mackelprang, R., Myrold, D. D., Jumpponen, A., Tringe, S. G., Holman, E., Mavromatis, K. and Jansson, J. K. (2014) ‘FOAM (Functional Ontology Assignments for Metagenomes): a Hidden Markov Model (HMM) database with environmental focus.’, Nucleic acids research, 42(19), pp. 1–7. doi: 10.1093/nar/gku702.

Ratel, D., Ravanat, J.-L., Berger, F. and Wion, D. (2006) ‘N6-methyladenine: the other methylated base of DNA.’, BioEssays: news and reviews in molecular, cellular and developmental biology, 28(3), pp. 309–15. doi: 10.1002/bies.20342.

Reisenauer, A. and Shapiro, L. (2002) ‘DNA methylation affects the cell cycle transcription of the CtrA global regulator in [i]Caulobacter[/i].’, The EMBO Journal, 21(18), pp. 4969–4977. doi: 10.1093/emboj/cdf490.

Russell, J. A., Zayas-Leon, R., Wrighton, K. and Biddle, J. F. (2016) ‘Deep subsurface life from North Pond: enrichment, isolation, characterization and genomes of hetertrophic bacteria’, Frontiers in Microbiology, 7(May), pp. 1–13. doi: 10.3389/fmicb.2016.00678.

Seitz, K. W., Lazar, C. S., Hinrichs, K., Teske, A. P. and Baker, B. J. (2016) ‘Genomic reconstruction of a novel, deeply branched sediment archaeal phylum with pathways for acetogenesis and sulfur reduction’, The ISME Journal. Nature Publishing Group, 1(10), pp. 1–10. doi: 10.1038/ismej.2015.233.

Souza, C. P., Almeida, B. C., Colwell, R. R. and Rivera, I. N. G. (2011) ‘The Importance of Chitin in the Marine Environment’, Marine Biotechnology, 13(5), pp. 823–830. doi: 10.1007/s10126-011-9388-1.

Spanevello, M. D. and Patel, B. K. C. (2015) ‘Thermoanaerobacter’, in Bergey’s Manual of Systematics of Archaea and Bacteria. John Wiley & Sons, pp. 1–6.

Srikhanta, Y., Dowideit, S., Edwards, J., Falsetta, M., Wu, H.-J., Harrison, O., Fox, K., Seib, K., Maguire, T., Wang, A.-J., Maiden, M., Grimmond, S., Apicella, M. and Jennings, M. (2009) ‘Phasevarions mediate random switching of gene expression in pathogenic Neisseria’, PLOS Pathology, 5.

Srikhanta, Y. N., Maguire, T. L., Stacey, K. J. and Grimmond, S. M. (2005) ‘The Phasevarion: A Genetic System Controlling Coordinated, Random Switching of Expression of Multiple Genes’, Proceedings of the National Academy of Sciences, 102(15), pp. 5547–5551.

Tolba, S. T. M., Nagwa, A. A. A. and Hatem, D. (2013) ‘Molecular characterization of rare actinomycetes using 16S rRNA-RFLP’, African Journal of Biological Sciences, 9(March), pp. 185–197.

Whitman, W. B., Coleman, D. C., Wiebe, W. J. and Colemant, D. C. (1998) ‘Prokaryotes: The Unseen Majority’, Proceedings of the National Academy of Sciences of the United States of America, 95(12), pp. 6578–6583.

Wion, D. and Casadesus, J. (2006) ‘N 6 -methyl-adenine: an epigenetic signal for DNA – protein interactions’, Nature Reviews Microbiology, 4(March), pp. 183–193. doi: 10.1038/nrmicro1350.

Wood, D. E. and Salzberg, S. L. (2014) ‘Kraken: ultrafast metagenomic sequence classification using exact alignments.’, Genome biology, 15(3), p. R46. doi: 10.1186/gb-2014-15-3-r46.

